# MASCARA: coexpression analysis in data from designed experiments

**DOI:** 10.1101/2024.02.29.582876

**Authors:** Fred T.G. White, Anna Heintz-Buschart, Lemeng Dong, Harro J. Bouwmeester, Johan A. Westerhuis, Age K. Smilde

## Abstract

Experiments in plant transcriptomics are usually designed to induce variation in a pathway of interest. Harsh experimental conditions can cause widespread transcriptional changes between groups. Discovering coexpression within a pathway of interest (here the strigolactone pathway) in this context is hampered by the dominant variance induced by the design. Minor changes in experimental conditions not controlled for may affect the plants, leading to small coordinated differences in genes within pathways of interest and related pathways between replicate plants in the same controlled experimental condition. These systematic differences are usually averaged out, but we argue here that they can be used to improve the detection of genes that co-express. We introduce a novel framework “MASCARA” which combines ANOVA simultaneous component analysis and partial least squares to remove the experimentally induced variance and investigate multivariate relationships in the non-designed variance. MASCARA is tested against a selection of competitors on simulated data, created to mimic a designed transcriptome study, where its benefit is demonstrated. In a coexpression analysis of a real dataset MASCARA detects several uncharacterised but relevant transcripts. Our results indicate that there is sufficient structure left in a typical dataset after correcting for experimental variance and that this residual information is useful to investigate coexpression.

**Author Summary:** Experiments in the life sciences usually purposefully induce significant variance between different treatments, in order to activate or repress certain mechanisms of interest. Whilst this is necessary it can make it challenging to detect meaningful relationships within pathways of interest, particularly when the experimental conditions are drastically different. Instead of focusing on the drastic changes in response due to the different treatment, MASCARA uses the systematic synchronous variances between replicates to find related features within the pathway of interest. Through simulation studies and application to a real dataset, we demonstrate the effectiveness of MASCARA in detecting relevant transcripts and extracting coexpression patterns from gene expression data.

## Introduction

Plant transcriptome studies typically involve the use of designed experiments which aim to induce variation in the expression of genes of interest by controlling one or more experimental factors. This can be done through, for example, varying the level of certain essential nutrients, temperature, water availability or knocking out or overexpressing a certain gene. Often these transcriptome data are used to identify genes of interest based on their expression profile, for example in biosynthetic pathways.

Analysis of RNAseq data generally adheres to one of two approaches: differential expression analysis (DE; (1); (2)) or coexpression analysis (CoE; (3); (4)). DE is used to determine which genes are expressed differently between two or more experimental conditions, highlighting known or novel pathways affected by the experimental conditions. CoE, on the other hand, aims at discovering genes that are part of an already partially characterized POI. Here, the expression profiles of known genes of the POI (termed “baits”) are used to detect novel pathway members with similar expression patterns (here termed “targets”).

Figure 1 outlines pathway nomenclature that will be used throughout this text. Panel *A* represents hypothetical pathways including a pathway of interest (POI), related pathway (RP) and an unrelated pathway (URP). All three types of pathway are assumed to be affected by the experimental design. URPs are defined to be pathways not directly interacting with the POI. In the hypothetical example displayed in Figure 1, taking the purple triangles as baits, the desired outcome of a coexpression analysis would be to detect all other purple and orange coloured genes and not the light green genes. However, these connections are not always preferentially detected, this is due to the presence of different sources of variance in the data.

**Figure 1:**
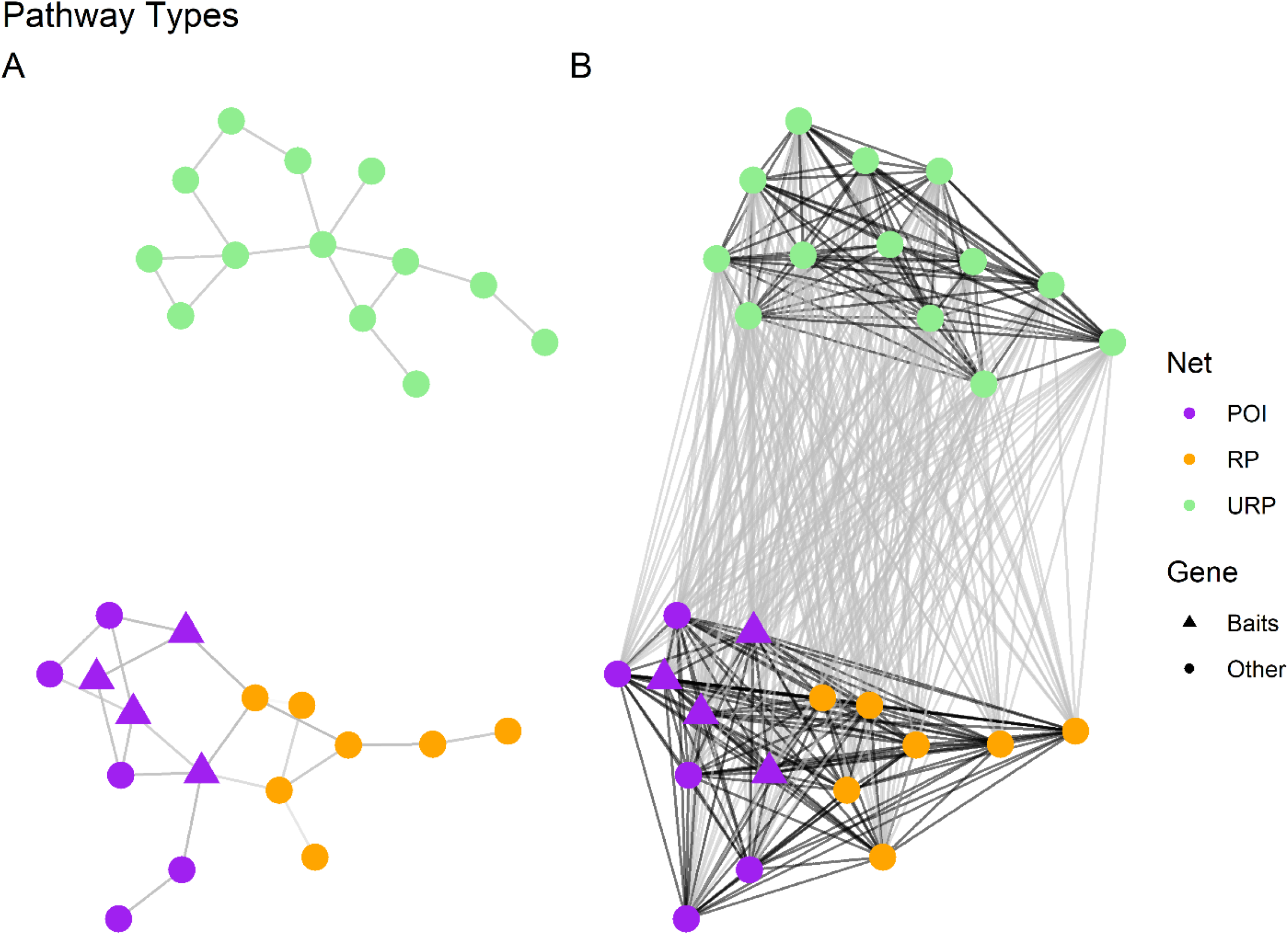
Pathway types exemplified, POI; pathway of interest, RP; related pathway, URP; unrelated pathway. Panel A represents hypothetical biological pathways that are all affected by the experimental design and thus all contain DE genes, here the edges represent real connections between the nodes/genes. Panel B highlights the problem with detecting these pathways when dealing with data from designed experiments, here edges represent putative connections defined by coexpression analysis; black is desired (within network detection), grey is non-desired.

When dealing with experimentally designed data, we have to consider two key sources of variance: between-group and within-group. Between-group variance pertains to the differences between experimental groups (or conditions). This type of variance is of interest when conducting DE studies. Within-group variance, on the other hand, is the variance within experimental groups (among replicates), this is not interesting for DE studies but may contain information useful to CoE. It is the excessive between-group variance that leads to the densely connected nature of panel *B* from 1. Concretising this with a biological example; in (5) a phosphate (P) starvation was applied to rice plants to induce strigolactone (SL) biosynthesis, to study the change in gene expression in comparison to control plants with normal supplies of P. The SL pathway is known to be triggered through P deficiency. However, as P is one of three key nutrients for plant growth, deficiency of this nutrient causes widespread differential expression throughout the transcriptome ((6)).

Figure 2 exemplifies between-group and within-group driven correlations from a real dataset ((5)). The plots show the relationship between two genes of the POI (*A* and *C*) and the relationship between two genes that are not part of the same pathway (*B* and *D*). The total correlations in both *A* and *B* are high but correlation in *B* is driven by the between-group variation and not by correlations within experimental groups. The within-group correlation is lower than the total in both cases, but there is a clear positive linear relationship in *C*: P- condition in red. The within group correlation (r) is higher for the first transcript pair (*C*) than the second (*D*), which makes it possible to discern between a POI gene or an URP gene. Hence, we need to distinguish *total CoE* which is calculated on the total variance and *within-group CoE* where the focus is on the shared variance within experimental groups.

**Figure 2:**
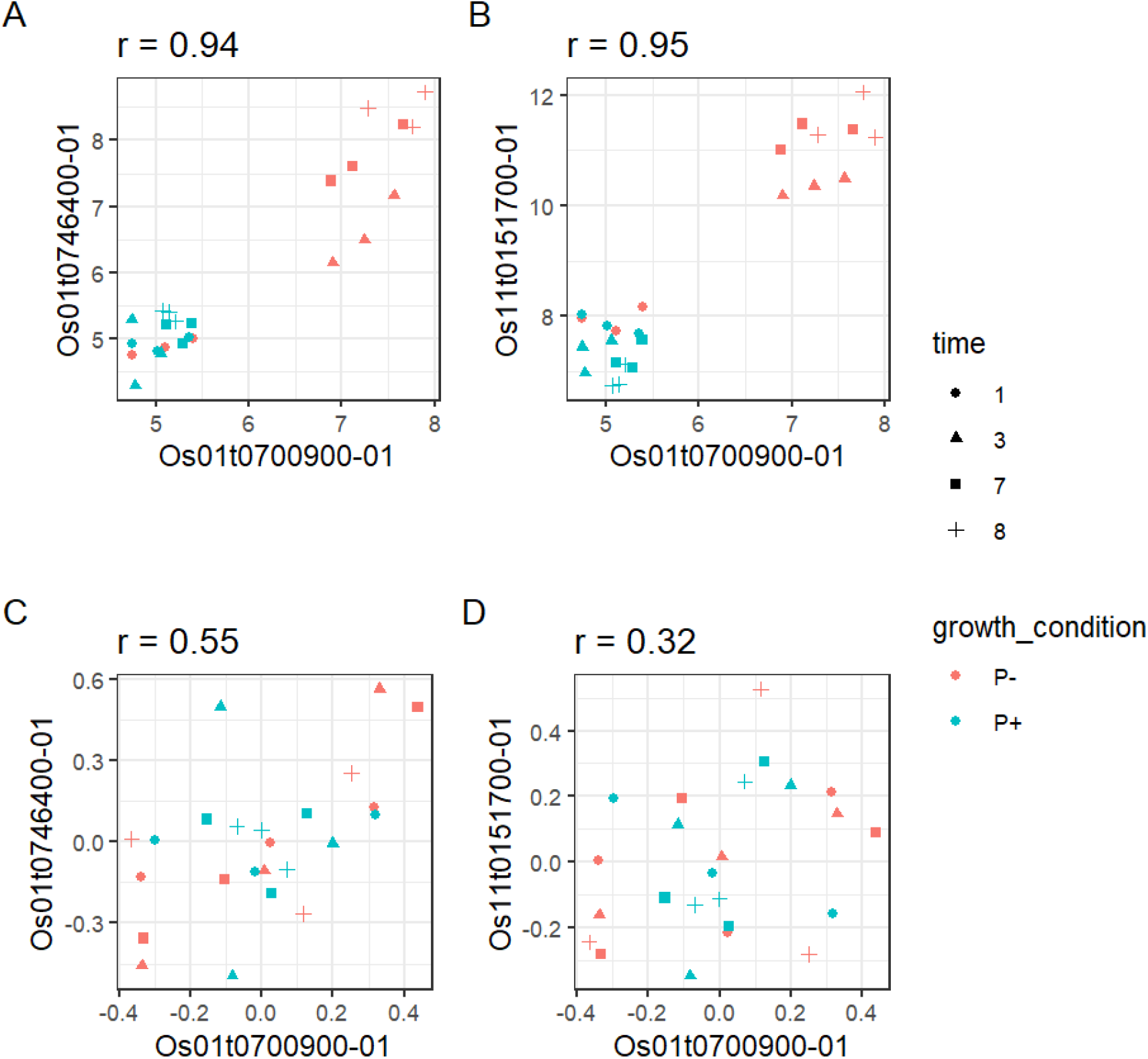
Correlation examples, Pearson’s correlation coefficient (r) indicated in facet titles. Data from (5). X axes, expression levels of MAX1 gene (Os01t0700900–01) in rice, Y axes expression levels of example coexpression candidates, aribitrary units. Top row A-B; total CoE, bottom row C-D within-group CoE from variance partitioning the data in A-B. Gene pairs in both A and B have strong total correlations. Correlation in A is also exhibited within a group, B has strong total correlations mainly due to the differences between groups. These examples are indistinguishable from eachother without variance partitioning.

From a plant biology point of view, the between-group (across treatments) and the within-group (across replicates) variance can be quite different. However, both may contain valuable biological information. The between-group variance is in principle larger because groups are the result of treatments that are usually selected because they affect the POI. Within-group variance (variance across replicates), if too large, can be an issue in DE analysis, we argue that in co-expression analysis the information present in this variance can be valuable. Variance between replicates is caused by non-controlled variations in plant development and environmental conditions. One replicate plant within a group may be slightly bigger than the others and therefore respond slightly different to a treatment.

This could result in slight differences in bait genes between the replicates, but also the genes that we want to identify with a co-expression analysis. There may be many other factors that are inducing this within-group variance, but it is quite likely that if a factor induces a change in the treatment response in one replicate, that this response difference occurs throughout the entire process or pathway in that one replicate. This means that the within-group variance likely contains valuable information and by averaging out this variance, as is done in differential expression analysis, we are losing information. Not making the distinction between types of variance results in *total CoE* associations driven by the differences between treatment groups as opposed to within group correlations. Haider and colleagues (5) used ranked Pearson correlations to determine (total) coexpression candidates. Due to such a strong effect of the experiment, many genes were affected on top of the desired SL genes. This masked the within-group correlation between the baits and the targets (remaining unknown SL genes). This problematic issue was also highlighted recently by (7) where group sampling assumptions are considered, concluding that between/within variance sources must be considered when investigating correlations.

Hence, both from a data analysis as well as from a biological perspective it is relevant to make a distinction between the different types of CoE. The focus of this paper will be on developing methods to investigate within-group CoE.

For a successful coexpression analysis it is necessary to have a predefined set of baits that are already known to be involved in the pathway under study. Methods like ranked correlations and partial least squares regression (PLS; (8); (9)) are able to investigate relationships between a set of baits and the other measured genes. In ranked correlations the relationships are quantified at a bivariate level. In PLS regression the relationships can be investigated in a multivariate way, in its simplest form we can see which genes are most predictive of the expression of one bait. This can be extended to a PLS2 model where relationships are investigated between a set of multiple baits and all other genes. These methods do not by default account for the different variance types discussed above. Hence, they are not directly suitable for within-group CoE.

The most widely known and applied method that investigates variance sources is the analysis of variance (ANOVA; (10)). ANOVA Simultaneous Component Analysis (ASCA; (11)) and its extension to imbalanced data via the generalised linear model (ASCA+; (12)) have been developed to combine Principal Component Analysis (PCA) with ANOVA and can therefore account for experimental design in dimension reduction. ASCA is of interest due to the variance partitioning. This is typically used to estimate the effect matrices that are calculated from the design and particularly the within-group variance structure which is maintained in the residuals of the model. Both PLS and ASCA have been used for DE ((13), (14), (15), (16)) and CoE ((17), (18)). ASCA and PLS have been combined before in (19) creating ANOVA-PLS, however the goal of their study was not coexpression analysis.

Neither of these methods are optimal by themselves for CoE in the context of large between group variance with known baits.

The baits are assumed to be part of the same POI and the remaining uncharacterised targets (POI or RP genes) are assumed to be numerous. Therefore, both the baits and the remainder of the data could be reduced to linear combinations in order to investigate relationships between a set of baits and putative targets. We introduce a new data analysis method called multivariate ASCA residual analysis (MASCARA). MASCARA is motivated by four key factors; (1) the presence of dominant between-group variance caused by the setup of the experiment, as well as, (2) ambient variance (structured effects caused by undocumented environmental factors i.e. temperature, light intensity or soil water content) within the replicates of each experimental group, (3) the need to investigate multivariate relationships and (4) the low number of samples. The method capitalises on ASCA to account for an experimental design and the multivariate analysis capability of PLS2 to find uncharacterised POI genes using a set of baits. Instead of focusing on the variance induced by the experimental design, as is a typical use of ASCA, this variance is removed, and the remaining residual variance is further analysed to investigate the multivariate relationships between baits and the putative targets in terms of within-group CoE.

To explore the utility of MASCARA, the method will be compared to ASCA, PLS2 and correlation analysis illustrated using an example RNAseq dataset (from (5)). Furthermore, this dataset is used as a guide to several simulation studies to validate the development of a novel approach to coexpression analysis. For these simulations we emulate both different levels of between- and within-group variance. We test the effects of size of between group differences, number of URP differential genes, measurement noise (random unstructured technical variance), structured ambient variance and number of replicates. The following sections seek to investigate which statistical methods are best suited for POI elucidation when a set of baits is already known and illustrate the benefit of MASCARA. Especially in data coming from experiments designed for differential expression analysis.

## Materials and Methods

### RNAseq - Rice strigolactone dataset (5)

In order to guide the simulations we illustrate the desired characteristics with a real dataset. The real data are a subset of samples from a study in which the root transcriptome of rice plants is measured under two growth condition (P+/P-) and measured at four time points (1, 3, 7 and 8 days post treatment). At each combination of growth condition and time point, the root transcriptome of three new plants was harvested for RNA sequencing. In this dataset, the curated list of SL pathway genes (contained in Supplementary Table 1) shows a specific profile: low variance baseline expression across all time points in the P+ condition but activation and increasing expression over time with P starvation, data were generated and preprocessed to variance stabilised counts as per (5).

Figure 3 shows the gene expression levels of 9 strigolactone pathway genes along with 9 highly differential genes that are induced by P deficiency, but are not part of the SL pathway. While the expression is low in P+ condition a clear increasing time profile can be observed in the P- condition, indicating an interaction between the growth condition and the time factors.

**Figure 3:**
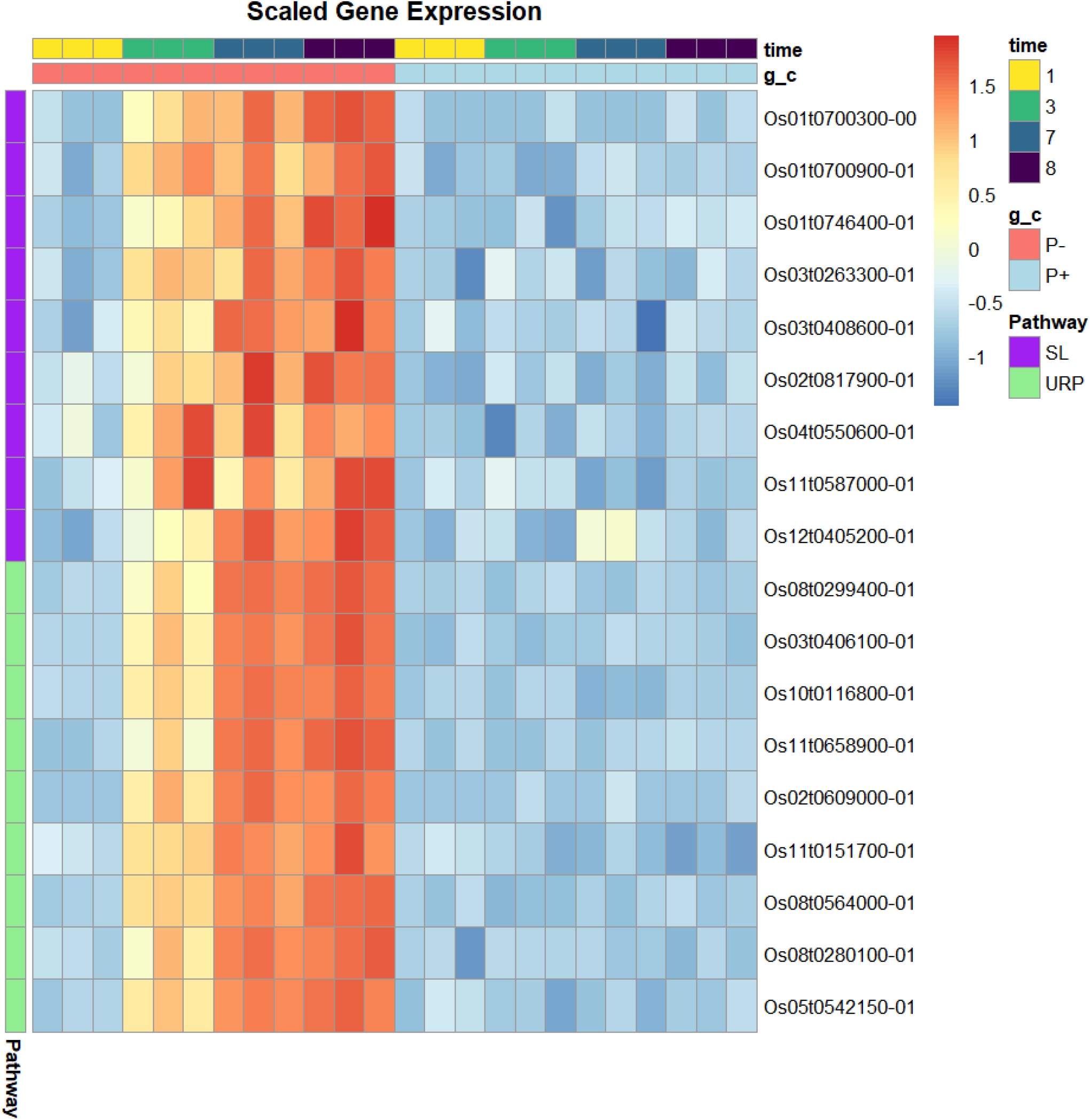
Real data overview. RNAseq of rice root. Heatmap of SL pathway and example highly differential genes. Data and preprocessing from (5), genes autoscaled i.e. blue indicates no/low expression and red indicates higher expression. IDs with functional annotations in supplementary table 1.

What is notable here is the replicate specific patterns that are apparent amongst the SL genes and largely absent between the SLs (purple) and the non-SL differential genes (green). For example, at time point 3, most SL genes are expressed lowest in replicate 1 and highest in replicate 3, while at time point 7 replicate 2 has a higher SL gene expression.

These within-group patterns are either not shared at all between the SL genes and the other DE genes or to a much lesser extent. This is due to the other genes not directly being part of the same pathway. There are of course other pathways and thus there are still some subtle replicate patterns in the non-SL genes but this effect is much clearer within our pathway of interest. Figure 4 shows the output of the ASCA decomposition in which the PCA scores and loadings of the combined effect of the Growth condition factor and the interaction of Growth condition and Time is explored. The score plot shows that the replicates of each condition are clustered while the difference between the conditions is much larger, as expected. While the 9 SL transcripts have higher loadings on the 1st PC, there are many non-SL transcripts with similarly (or more) extreme loadings. Thus many differentially expressed genes are found important that are not part of the SL pathway. In other words, ASCA does not specifically pinpoint SL genes (POIs) among others that are similarly or even more differentially expressed but not part of the SL pathway (URPs).

**Figure 4:**
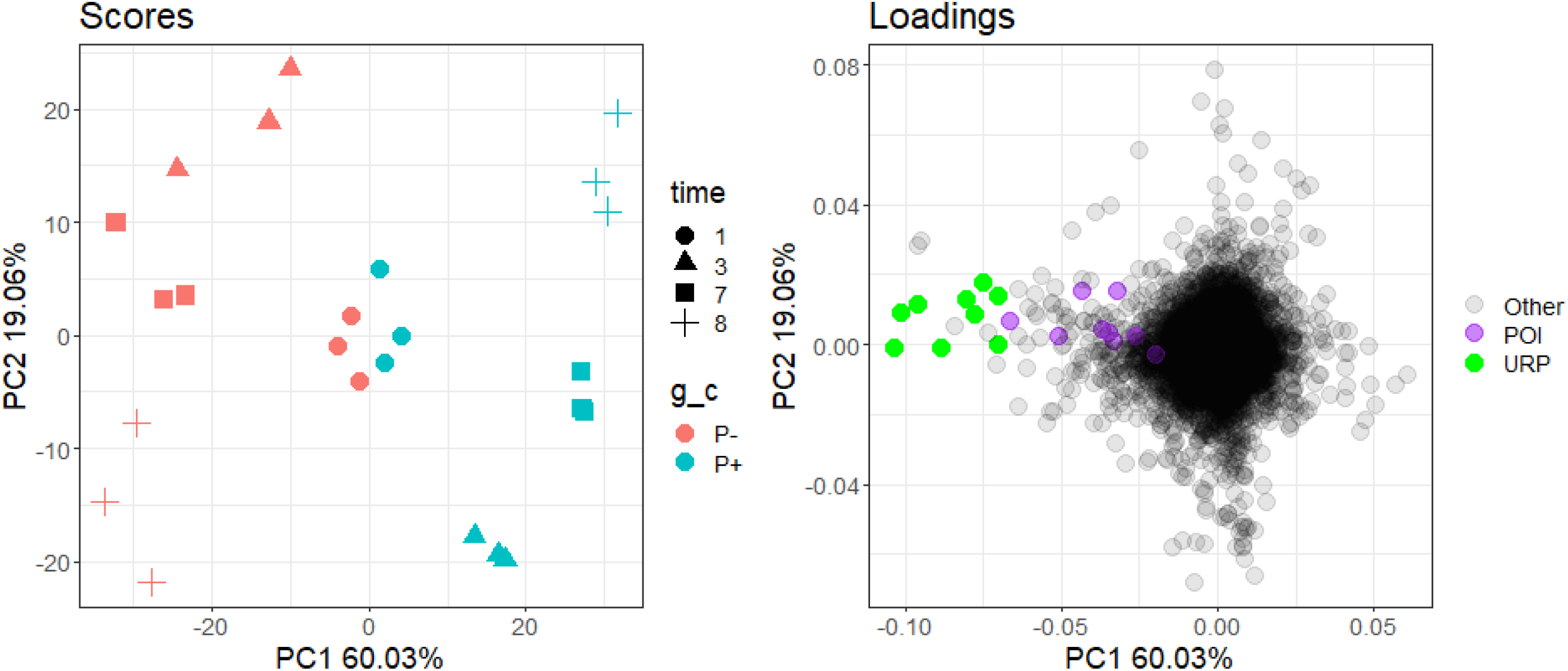
Scores and loadings of the ASCA model of the combined effect (condition + condition:time interaction). Scores: g_c indicates growth condition, time indicates days after starvation induction. Loadings: POI; SL genes and URP; other highly differential genes (indicated in purple and green respectively) show upregulation in the P- condition compared to P+.

This marks the goal of this paper, which is to detect within-group coexpressed genes (POI/RP) with a set of baits, preferentially to other differentially expressed genes that are not part of the same pathway (URPs, with a different within-group structure).

### Simulations

The problem in the real dataset is that the nutrient stress applied causes a major effect on many other genes besides our baits and targets. This widespread transcriptional regulation can mask the coexpression patterns of interest due to the large number of genes that are upregulated under the stress condition, making it more difficult to detect genes that are responding to a lesser extent, for example the SL pathway.

The simulations are built upon the results of the ASCA analysis of the combined effect of growth condition (g_c) and the interaction of growth condition and time. As can be seen in Figure 4 there is clear separation between samples due to the experimental factors growth condition and time. The variation between the replicates is only small. The green loadings of URP genes are strongly related to the experimental variation due to g_c. The simulated data is created by simulating experimental variation consisting of the main time effect *β* and the combined effect (*α*+ *αβ*), and the Ambient/Residual variation which exists of the structured ambient variance (**S**) and residual variation (**E**) representing measurement error, modeled with normally distributed random values *e*_*ij*_ = 𝒩(0,σ^2^).

Each of the sources of variation except the residual variation is simulated as low dimensional variation created from 2 scores (**T**) and loadings (**P**). For the experimental variation, the scores matrices **T**_*α*+*αβ*_ and **T**_*β*_ each consist of two orthogonal score vectors and their relative levels are indicated by the colour. The different colours in Figure 5 represent that the scores are orthogonal to each other. Each small block consists of the same score value for each of the replicates within that condition. For the Ambient variation, the scores **T**_*s*_ are different for each replicate plant. The values in **T**_*s*_ are obtained from a normal distribution with mean 0 and variance 1, i.e.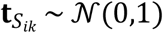. The **T** matrices represent the simulated scores for their respective factors. For blues, reds, greys and golds we have 4 different shades representing 4 unique values per factor level (total 8 unique values where reds and golds are sign flipped blues and greys respectively). Similarly oranges and pinks represent a growth condition independent time profile with four unique values.

**Figure 5:**
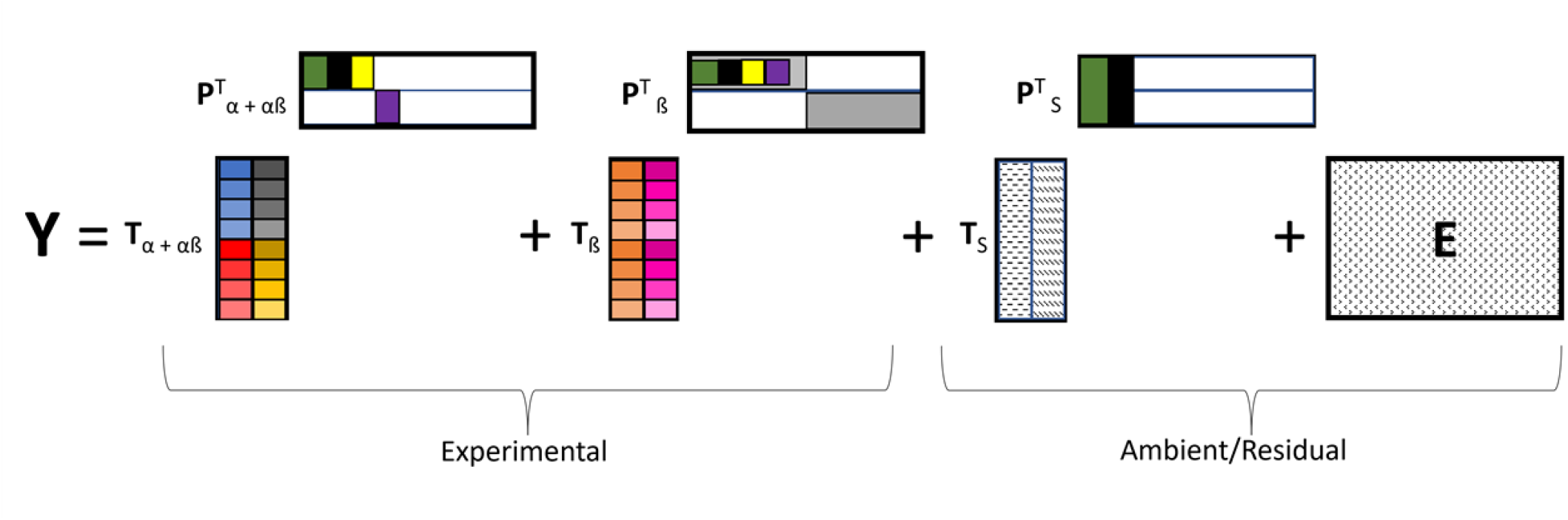
Coexpression simulation, simplified overview. Colours correspond to the same genes in the different loading vectors (**P**). Black; baits (known POI genes), green; targets (unknown POI genes), yellow; DE URP genes, purple DE URP genes with a different profile. In the score vectors (**T**); Blues, reds, greys and golds represent experimental conditions, oranges and pinks; time. Each of the textures in the ambient/residual part represent a distinct normally distributed set of random values.

In the loading matrices **P**, the different coloured blocks indicate different types of genes. The green block represents the bait genes of POI. The black genes represent “unknown” target genes from POI or RP. Yellow and purple represent genes from URP that are affected by the experimental variation in different ways. The grey blocks represent very small background variation for those genes, while the white blocks represent that those genes are not affected by that source of background variation.

In 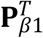 the baits, targets and other DE genes (all genes from POI, RP and URP) along with the remaining first half have similar positive values (represented by light grey) where the other 1000 have positive values in 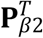 (grey), these are created to emulate the reality of the real data in which many genes are slightly correlated with eachother through the course of development, as well as to account for some nonlinearities which are otherwise ignored by our model. The remaining elements of 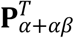 and 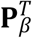 are populated with zeroes. **P**_*s*_ only has values for the baits (green) and target (black) genes. The loading values for the baits and targets are obtained as 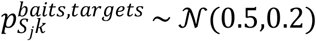, while the loadings for the other genes are 0.

To finalise, the response values all have an intercept 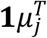. Similarly to the real data, the gene expression values for each experimental condition are simulated with three independent replicates per time-condition-combination. Each simulated data Y set thus consists of 24 samples and 2000 genes.

### Data simulation 1: Combined effect size and number of differential genes

The first simulation aims to determine how well the methods perform to find coexpressed genes in the presence of a dominant between-group variance structure and increasing numbers of differential URP genes (represented by the yellow block in Figure 5) that only have high between-group variance but no within-group variance.

In this simulation we have a set of 4 baits, 12 target genes and a variable number URP genes. The issue to address is the presence of many differential URPs; those that have strong between group correlation but no within group correlation with the POI. We incrementally increase the number of genes in the yellow block and decrease number of white genes with 0 loadings. As well as the number of DE URPs we also independently control the size of the experimental variance through the parameter *d* in Equation in (1). This approach controls the size of the experimental variance as well as the number of genes responding while keeping a fixed amount of ambient or within group variation.

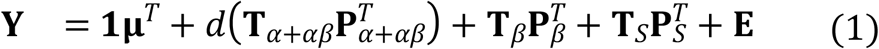

### Data simulation 2: Replication

The second simulation focuses on the effect of a different number of replicates. We investigate a range of values for sizes of between experimental variance (*d*) and random noise variation (*l*) across random structures (which correspond to 20 repeated simulations of randomised values for all **T** matrices).

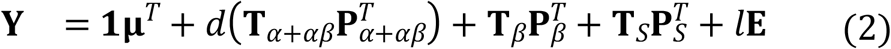

On these 20 datasets, per condition, the replicate number *n*_*r*_ was also varied: the number of replicates *n*_*r*_ ∈ 3:15 per experimental condition were sampled from a maximum *n*_*r*_ of 15.

With this setup we test the feasibility range of MASCARA with respect to the ratio of 3 key variance parameters namely: 1. Within group correlation structure 2. Between group correlation structure 3. Random noise

Here we set a fixed size of within group correlation structure 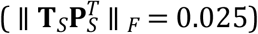 as well as number of non-pathway DE genes to 30 (yellow bar size Figure 5) across all iterations of this test. We vary the sizes of the other two variance parameters (*d* and *l*). Furthermore we also show the effect of replicate number and demonstrate that MASCARA (along with the other methods) can be applied to data with generic profiles, not limited to the specific structures that were used to demonstrate the differences in coexpression analysis with simulation 1. Here we create 20 datasets with random combined effect and within group correlation structures.

### Selection of genes of interest

#### Ranked correlations

Pearson’s correlation coefficient 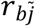 is calculated between all bait genes *b* = 1‥*B* and all other genes 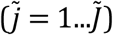. Ranks 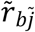 are assigned for each 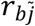 such that the highest correlation for each bait gene *b* receives the lowest rank. These ranks are averaged across all baits *B* (see supplementary equation (14)).

#### ASCA+

ANOVA simultaneous component analysis ((11)) involves splitting data into effect matrices which are defined by the experimental design structure, after which a PCA is applied to the effect matrices or combination thereof. In ASCA+ this variance partitioning is achieved through the generalised linear model as described in (12). ASCA+ addresses the issue of biased effects matrices in unbalanced designs. The following is a summary of the algorithm description provided in (12).

The multivariate generalised linear model (GLM) can be expressed as:

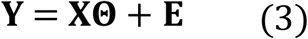

where **Y** is the response matrix of dimension *I* × *J*, **X** is the model matrix of dimension *I* × *p*, containing the sumcoded indicator levels for each of the levels for each factor and their interactions, **Θ** is the parameter matrix of dimension *p* × *J* and **E** is the error matrix of dimension *I* × *J*.

Through Ordinary Least Squares (OLS) we can obtain the (unbiased) estimators of the model parameters:

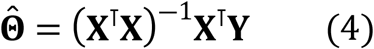

The effect matrices for different terms of the model can be obtained as:

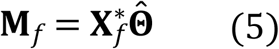

where: **M**_*f*_ is the effect matrix corresponding to effect *f*, 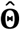 is the matrix of estimated parameters obtained from OLS and 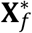 is a new model matrix obtained by keeping in **X** only the block **X**_*f*_ and replacing all other columns with zeros as per:

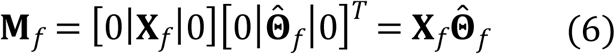

The above is achieved through the sum coded indicator matrix as illustrated further in (12). Furthermore, the matrix **E** is estimated as the residual matrix of the model:

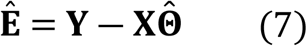

Throughout this text when referring to the calculation of ASCA/ASCA+ model on simulated or real data the following model is calculated:

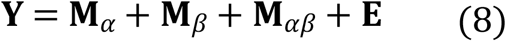

With genes selected from:

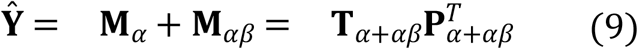

#### PLS regression

Partial Least Squares (PLS2) is a multivariate regression technique used for modeling relationships between two sets of variables. Here we split our data matrix **Y** into sub-matrices;

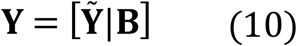

The response matrix **B** contains only the baits (relating to the green bar in the loadings of Figure 5) and the predictor matrix 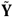 contains the rest of the data, this can then be expressed as:

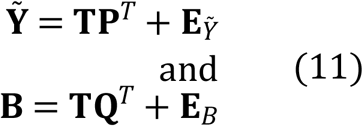

This model is estimated in our work using the SIMPLS algorithm ((20)) as follows:

1. For the 1^*st*^ component:
  a. Compute the left singular vector **u**_1_ 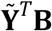
  b. 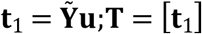
  c. 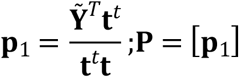
2. For the following components *k* = 2…*K*:
  a. Compute the left singular vector **u**_*k*_ of 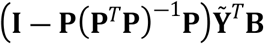
  b. 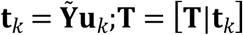
  c. 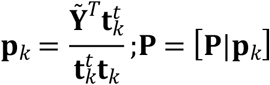

Under the conditions of orthogonal scores; 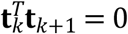 and normalised weights; 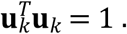. Loadings for **B** can be obtained with a least squares step **Q** = (**T**^*T*^**T**)^―1^**T**^*T*^**B**.

In the following simulations and application the number of components calculated by PLS2 is set to 2 to create an interpretable model.

### Variable importance in projection (VIP)

In this work genes are selected in both ASCA+ and PLS through VIP which is a way to measure the contributions of each variable to the underlying models. According to:

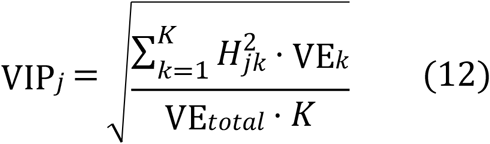

Where *k* = 1…*K* represents the number of components in the respective models, *j* indexes a gene, *H* is substituted for the loadings 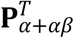 or weights **U** (for ASCA+ and PLS respectively) and VE is the variance explained, where *VE*_*k*_ is the amount of variance explained by component k and *VE*_*total*_ is the variance explained by the whole model. In ASCA+ we calculate the VIP scores in the latent space of **Y**_*α*+*αβ*_ as described in (12). In PLS we use the VE from the latent space of **B** using the weights (**U**) from the latent space of the predictor matrix **Y**.

### Multivariate ASCA residual analysis (MASCARA)

MASCARA combines the effect estimation with GLM (from ASCA+) with PLS2. The method entails estimating effect matrices with the GLM step of ASCA+ as described above to obtain the estimated residual matrix **Ê**. The variance explained by all effect matrices of the experimental factors and interactions is removed. What remains in this **Ê** matrix is individual variance of each replicate that is assumed to contain not just random technical variance but some form of structured variance induced by ambient (uncontrolled) effects. The matrix **Ê** from equation (7) is substituted as **Y** into PLS2 after being split into baits and remainder as per equation (10).

The genes are ranked based on target projection ((21)) of the loadings **P** onto the mean vector of the bait loadings **Q** in 2 components: 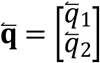 where 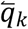 is the mean of the *k*^*th*^ component of the PLS2 model. The target projected loadings **P**_*TP*_ are calculated such that 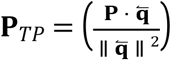. Higher values in the target projection equate to stronger positive associations with the center of the baits. Large negative values can also be interesting candidates for strong negative associations, although only positive associations are considered here. This approach allows a general direction in the PLS space to be defined by multiple baits and assumes that these baits are highly correlated with one another.

### Performance evaluation; Log2 geometric mean rank

To evaluate performance we use *log*_2_ geometric mean rank 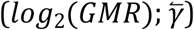, defined as:

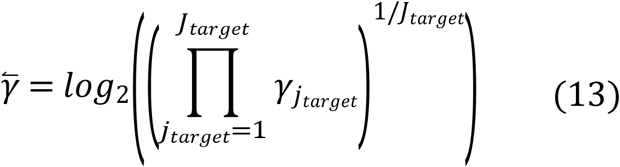

for *J*_*target*_ targets. A lower 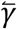 indicates better performance.

## Results

### Simulation 1: Combined effect size and number of differential genes

For this simulation we have some experimental variance for the combined effect and time 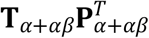 and 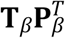 respectively) a fixed amount of random noise **E** and a fixed amount of structured ambient/residual variance 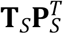, within which our 4 baits and 12 targets share a correlation structure. This reflects the situation expected from the real data example - drastic effect caused by nutrient deficiency, which also activates the SL pathway. The POI genes are expected to share some variance that is independent to the experiment, the 4 baits and 12 targets load on to the structured part of the noise 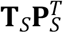.

We can see in Figure 6 that MASCARA was neither affected by the presence of an increasing number of non-pathway DE genes nor the size of this effect. Correlations, ASCA and PLS, however, all showed the expected breakdown in performance due to the injection of more DE URP genes. This was the expected result as the experimental between group variance is removed with MASCARA.

**Figure 6:**
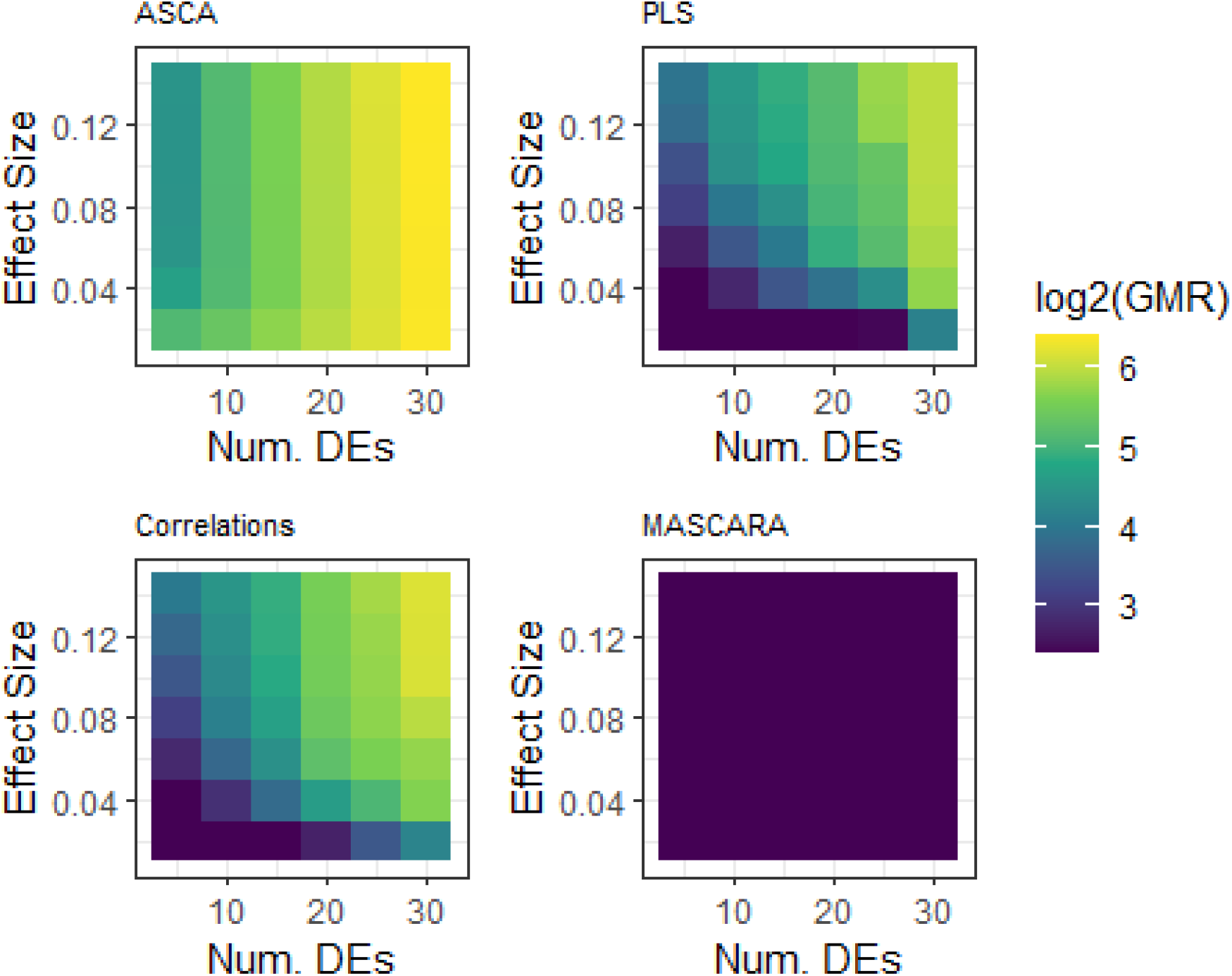
Results of simulation 1 for all 4 methods. Log2 geometric mean rank (log2(GMR)) as a function of the number of differentially expressed URP genes (x-axis) and combined effect size (y axis); 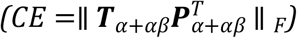, lower log2(GMR); dark blue, indicates better performance. ASCA model calculated per equation (8), PLS model per equation (11).

**Figure 7:**
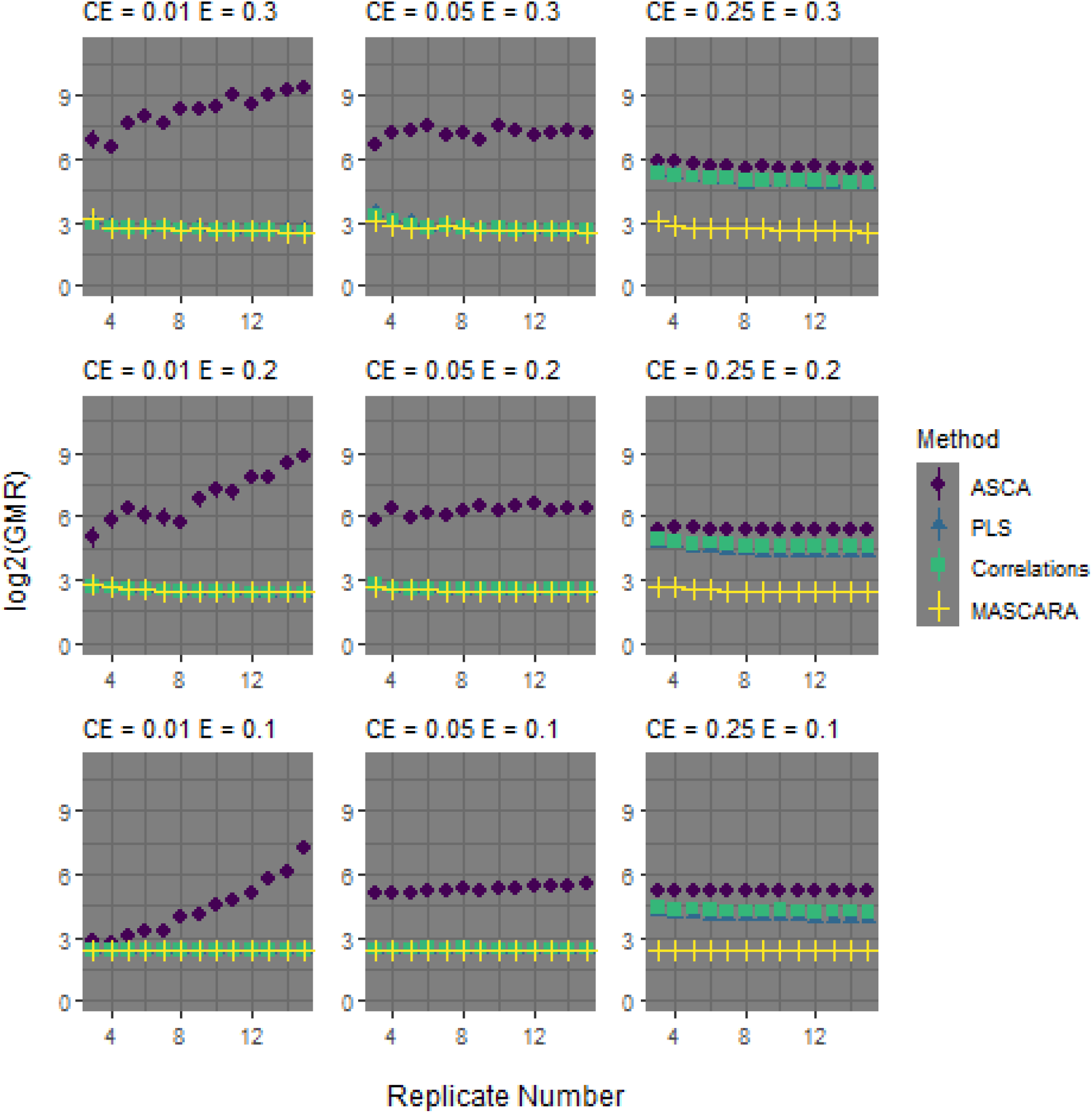
Simulation 2 results. Replications and variance type feasibility ranges. X axes; replicate number, y axes log2 transformed geometric mean rank. Each facet is created with set parameters for combined effect size 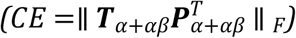 and random gaussian noise size (**E** =∥ **E** ∥ _F_). Dots are median performance across 20 random structures. ASCA model calculated per equation (8), PLS model per equation (11).

### Simulation 2: Replication

It was expected that all geometric mean ranks converge towards a minimum with increasing numbers of replicates, see 7. Perfect detection of 12 targets in top 12 candidates is achieved with a log2 geometric mean rank of 2.4, MASCARA was able to achieve this, however, the other methods were perturbed by the presence of the 30 differential URP genes. With more replicates MASCARA always outperforms competitors. With higher values of *d* (larger combined effect size) (CE) MASCARA outperforms the other methods but its performance is affected by amount of random noise (E) size indicated by *l*.

In simulation 2 we fixed the number of differential URP genes to 30, however this is less than 2% of the total number of genes in each dataset. In the real data example more than 10% of of the total number of genes are differentially expressed. As shown in simulation 1 (Figure 6) higher numbers of DE URP genes results in higher log2 geometric mean rank (log2(GMR)) of the targets for correlations, ASCA and PLS. MASCARA is designed specifically to mitigate the effect of large numbers of DE URP genes. Here we show that this is achieved.

### Real Data Applications

It is apparent that MASCARA has a particular use case for coexpression analysis in data where there is a lot of experimentally induced between-group variance across many genes as well as some within-group correlation structure shared between POI genes that is caused by independent non-controlled ambient factors.

Here we illustrate an application of MASCARA to the dataset from (5). For this we take 4 core strigolactone genes (Os11t0587000, Os04t0550600, Os01t0746400 and Os01t0700900 of the 9 used for illustrations in Figures 3 and 4) as baits and investigate their relationship with top candidate genes.

In the real data from (5) some plants were subjected to massive stress with the phosphate limitation. This is, therefore, reflected by large differences in the transcriptome compared to those with a normal supply of P. This limitation affected many more genes than those directly involved in SL biosynthesis. Furthermore, there appears to be a stronger relationship amongst the SL genes in the ambient variation compared to their correlations with other DE URP genes, as displayed in Figure 8 where the blue distribution indicates correlations between residuals of the 9 POI transcripts from Figures 3 and 4 and the red indicates correlations between residuals of these 9 and top DE URP genes.

**Figure 8:**
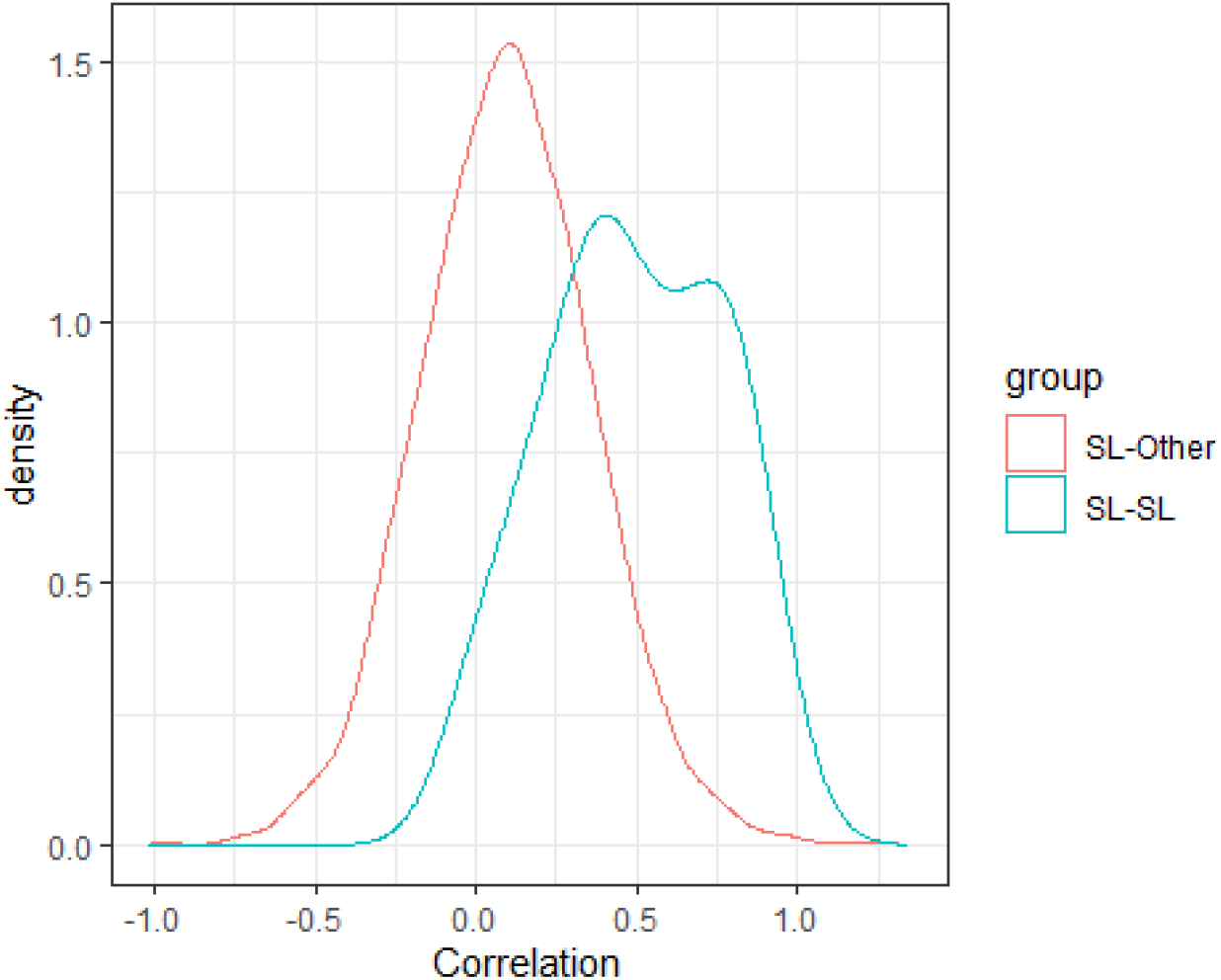
Distributions of Fisher transformed Pearson correlations (supplementary equation (15)) in residuals from the ASCA model on the data from (5). Blue indicates correlations between SL pathway genes, red indicates correlations between SL pathway genes and top 10% DE URP genes (upregulated in P- over P+).

Some example top coexpression candidate genes are shown in Figure 9. The problem of dominant between group variation is again exemplified here. Several of the top candidates detected by correlations (ASCA and PLS, results not shown) show no indication of within group correlations. Candidates selected with MASCARA, on the other hand, consistently show within group correlations with the baits. Furthermore, the strigolactone pathway, which is still under active investigation, has several uncharacterised biosynthetic steps. These are thought to be carried out by cytochrome p450s. The top candidates from MASCARA contain at least three (putative) cytochrome p450 genes that have not yet been biologically investigated in SL biosynthesis, as well as, a methyltransferase and several putative carotenoid synthesis genes. MASCARA also detected genes linked to Gibberellins (GA) preferentially to general phosphate starvation genes. The GA pathway has been linked to the strigolactones ((22), (23)). The interaction between these two pathways has been characterised recently under nitrogen limitation by (24).

**Figure 9:**
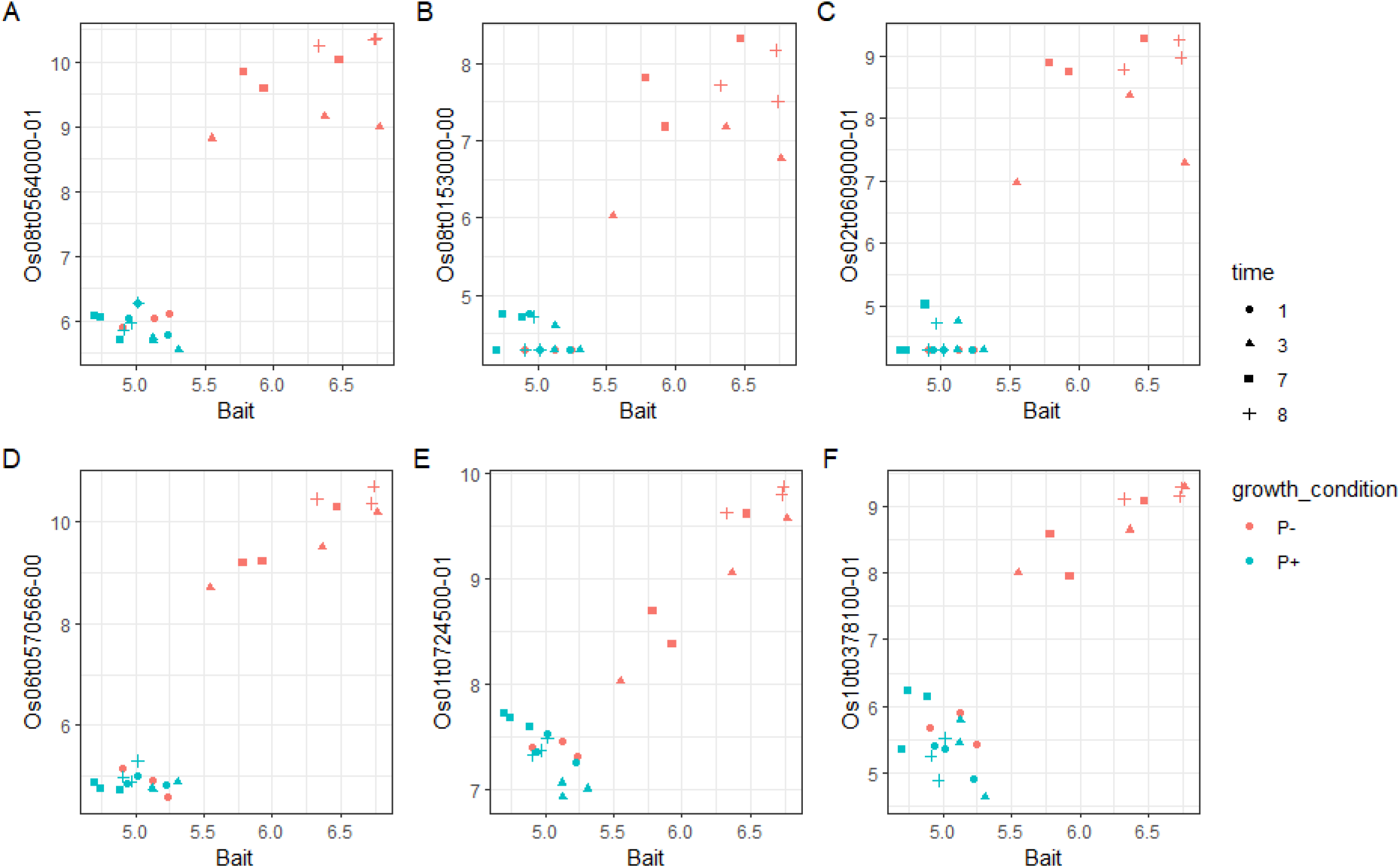
Example candidates from A-C ranked correlations and D-F MASCARA. X axes; transformed expression level of one of the bait genes (CCD8), y axes; expression levels of three example coexpression candidates for each method (see (5) for preprocessing details).

**Figure 10:**
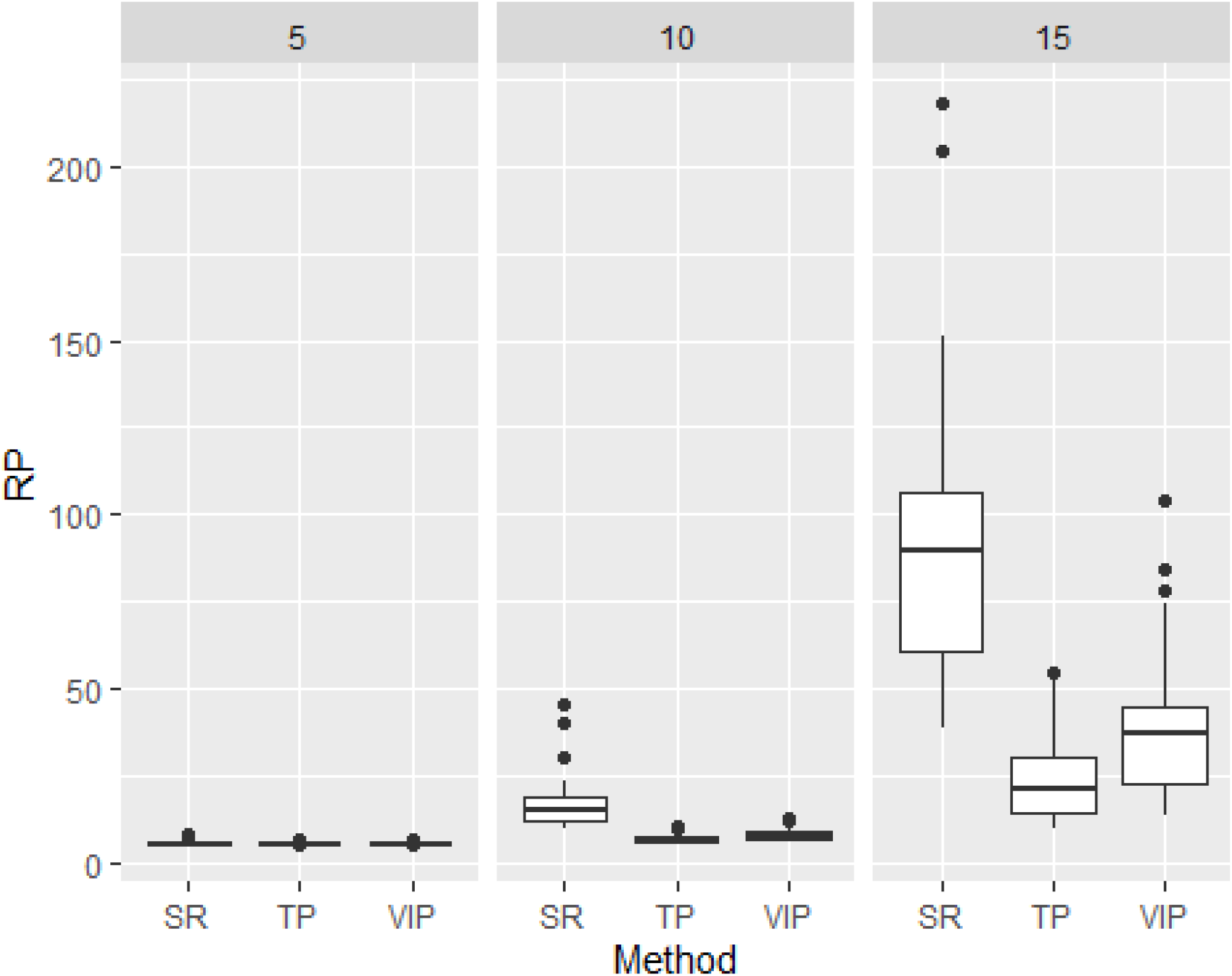
Feature selection simulation, selectivity ratio; SR, target projection; TP and variable importance in projection (VIP) compareed at 3 levels of noise, indicated in facet titles.

## Discussion and conclusions

In this article we introduced multivariate ASCA residual analysis (MASCARA) as a method for coexpression analysis in data containing a dominant variance structure from a known (experimental) source. MASCARA estimates effects through the GLM with a design matrix to remove the variance between conditions, then the residual matrix can be decomposed through PLS2, with feature ranking through target projections. This framework enables the investigation of multivariate relationships between a set of bait genes and all other genes in the dataset. MASCARA thrives on the fact that within the typical experimental setup small systematic variance exists between replicates due to variations in ambient factors that affect the plant, and therefore the variance structures in the resulting data.

Through simulation studies we showed the conditions under which MASCARA is suitable for coexpression analysis and compared it to a selection of other coexpression methods (ASCA+, PLS2 and correlations). These methods were applied to data from (5) exploring the strigolactone pathway and simulated data based on the structure of the Haider data. In a coexpression context, within certain assumed parameters of variance structure (dominant between-group, existing within-group), MASCARA outperformed competitors. Ranked correlations, ASCA ((11), (12)) and PLS ((9)) are all affected by dominant experimental between-group variance, which is a common occurrence in experimental data, exemplified here by the dataset from (5) but discussed elsewhere ((25), (26), (27)).

The feature selection stage of MASCARA uses an average vector constructed from the loadings of the baits onto which the loadings of other genes are projected. This approach is similar to target projection in (21) although we construct the target vector from averaging the loadings in a PLS2 model as opposed to already having it defined with the univariate response in PLS1. There are many established methods for feature selection in PLS which include Variable Importance in Projection (VIP) and Selectivity Ratio ((28)), however these methods focus on the importance of variables in the construction of the model rather than a directional association with the response block as with the 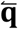 target projection stage. With this averaged target projection approach, as an extension of Kvalheim’s target projection in PLS1 models (21), we can use the information in a set of baits to define a direction in the latent space which can then be used as an axis to determine the relatedness of other genes. This assumes moderate to high correlation in the residuals of the baits, which is a realistic assumption in coexpression analysis although should be verified in the selection of baits.

Our analysis framework does not consider different correlation structures within the residual variance, this could be an issue especially in the cases similar to (29) where different combinations of experimental factors show different within-group correlation structures. This type of relationship can be seen in Figure 9 where the P- experiments show high positive correlation between the genes of the same pathway while the P+ experiments show a negative correlation. With MASCARA these will not be preferentially detected, as the focus is on detecting positively associating features. Ideally there would be enough samples to calculate robust correlations per group, to account for possible rewiring under different conditions, but in our case we focus on experiments where there are generally not enough replicates to do this. Experiments with higher replication number are always desirable from the point of view of data analysis. However, this is not always feasible due to time and space constraints. We tested the effects of replication number and found that MASCARA can outperform its competitors with lower replication, especially in data containing many differential genes between experimental conditions. This advantage holds as long as there is a realistic amount of structured variance caused by undocumented ambient factors.

In coexpression analysis, there is no need for class balance, it is in fact better to have more data points in the class where the pathway is induced (P- in the example dataset). When the POI is only active in the case condition then the control samples are mostly redundant. With more samples in the condition of interest the reliability of the coexpression analysis will improve. The MASCARA approach indicates that the typical experimental design used for DE is not optimal for coexpression studies. The fact that the residuals need to be isolated to find within group variance confirms the notion that coexpression analysis warrants different experiments than those used for DE studies. If quantification of differential expression is also needed, for example to confirm the pathway under study is affected by the treatment, but the main focus of research is coexpression, then it is an option to have just a few samples in a control condition to confirm the experimental treatment has caused the desired effect. This was not tested explicitly here, however if pathway genes are not active in a given condition then this set of samples does not provide any useful coexpression information.

This work has outlined and tested a basic use of MASCARA where the relationship between baits and unknown pathway genes is predefined to be positive. The method is not limited to the calculation of positive associations, however performance in detecting negative correlations (indicative of inhibitory or negative feedback processes) or more complex relationships has not been tested within the current simulations. Our approach was based on the structure of the study by Haider with RNAseq measurements of rice plants, however the method is applicable across other organisms and omics types. This holds under the assumption that the set of baits have a multivariate relationship between themselves and multiple other (unknown) genes. These genes are also related to the baits, as in our partially characterised strigolactone pathway example. In untargeted metabolomics datasets under similar experimental conditions these assumptions are also likely to be met.

Taken together our results indicate that the lack of ability to discern different types of variance in a dataset can compromise the detection of true coexpression patterns. MASCARA accounts for the different types of (experimental and ambient) variance and enables finer scale detection of within group variance structure. Code and data for this project are available at https://github.com/BiosystemsDataAnalysis/MASCARA.

## Acknowledgements

FW would like to thank Frans van der Kloet, Fentaw Abegaz, Roel van der Ploeg for statistical discussions, Imran Haider for consultation on the strigolactone pathway and the Data Science Centre of the University of Amsterdam.

## Supplementary

Target projection in PLS2 models has not been described in the literature, we therefore tested our approach against the most common methods for feature selection in PLS namely variable importance in projection (VIP) and selectivity ratio (SR). Here we have similar but reduced simulation set up to the replicate test section; 3 levels of unstructured noise; matrix F-norm sizes (as indicated in the panels). In each of the 3 tests the **T**_*s*_ is randomised 30 times resulting in different correlation structures between baits and targets. Target projection using 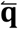 significantly outperforms the conventional approaches.

### Ranked correlations

Ranked correlations are calculated as:

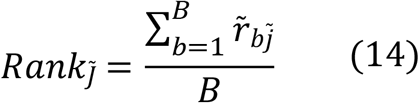

Where *B* is the number of baits and 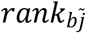 is the ranked Pearson correlation coefficient between gene j and bait b in the context of 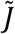 non-bait genes.

### Fisher transformed Pearson correlation coefficient

To calculate Fisher transformed Pearson correlations (z) with Pearson’s r:

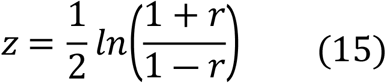

